# Effects of optogenetic stimulation of basal forebrain parvalbumin neurons on Alzheimer’s disease pathology

**DOI:** 10.1101/2020.04.26.062950

**Authors:** Caroline A Wilson, Sarah Fouda, Shuzo Sakata

## Abstract

Neuronal activity can modify Alzheimer’s disease pathology. Although overexcitation of neurons can facilitate disease progression, the induction of cortical gamma oscillations can reduce amyloid load and improve cognitive functions in mouse models. These beneficial effects of gamma oscillations can be caused by either optogenetic activation of cortical parvalbumin-positive (PV+) neurons or 40 Hz repetitive sensory stimuli. However, given the fact that cortical gamma oscillations can be induced by multiple mechanisms, it is still unclear whether other approaches to induce gamma oscillations can also be beneficial. Here we show that optogenetic activation of PV+ neurons in the basal forebrain (BF) increases amyloid burden, rather than reducing it. We applied 40 Hz optical stimulation in the BF of 5xFAD mice by expressing channelrhodopsin-2 (ChR2) in PV+ neurons. After one-hour induction of cortical gamma oscillations over three days, we observed the increase in the concentration of amyloid-β42 in the frontal cortical region, but not amyloid-β40. The density of amyloid plaques also increased in the medial prefrontal cortex and the septal nuclei, both of which are targets of BF PV+ neurons. These results suggest that effects of cortical gamma oscillations on Alzheimer’s disease pathology can be bidirectional depending on their induction mechanisms.

**Significance Statement:** Alzheimer’s disease (AD) is the most common cause of dementia. Although numerous molecular targets have been identified, the development of treatment is still a challenge. Accumulating evidence shows that artificial control of neuronal activity can modify AD pathology. In particular, the induction of cortical gamma (~40 Hz) oscillations can ameliorate AD pathology and improve cognitive functions. Here we show that optogenetic activation of parvalbumin-positive (PV+) neurons in the basal forebrain (BF) has opposite effects. By expressing channelrhodopsin-2 (ChR2) in PV+ neurons of an AD mouse model and optically stimulating BF PV+ neurons, we induced gamma oscillations and found increased amyloid burden. These results imply that AD pathology can be modified bidirectionally depending on induction mechanisms of gamma oscillations.

## Introduction

Amyloid plaques are a hallmark of Alzheimer’s disease (AD) along with tauopathy as well as neuroinflammation (Hardy and Selkoe, 2002; Herrup, 2015; Musiek and Holtzman, 2015; Hickman et al., 2018; Henstridge et al., 2019; Long and Holtzman, 2019). Although a wide range of molecular targets have been identified for pharmaceutical interventions, it is also important to develop alternative non-pharmaceutical intervention strategies given the complexity of AD pathology. One of such strategies is a neuromodulation approach (Iaccarino et al., 2016; Mirzadeh et al., 2016; Senova et al., 2018; Adaikkan and Tsai, 2020).

While abnormalities in neuronal activity have been associated with AD (Coben et al., 1983; Leuchter et al., 1987; Delbeuck et al., 2003; Jeong, 2004; Stam et al., 2007; Busche et al., 2008; Bero et al., 2011; Verret et al., 2012), recent optogenetic or chemogenetic studies in mouse models have demonstrated that the artificial manipulation of neuronal activity can modify AD pathology (Yamamoto et al., 2015; Iaccarino et al., 2016; Yuan and Grutzendler, 2016). For example, increased or decreased neural firing promotes or suppresses the spread of Amyloid-β (Aβ), respectively (Yamamoto et al., 2015; Yuan and Grutzendler, 2016). The induction of cortical gamma oscillations can reduce amyloid load and improve cognitive functions by activating microglia (Iaccarino et al., 2016; Adaikkan et al., 2019; Martorell et al., 2019; Adaikkan and Tsai, 2020).

Although this new approach based on gamma oscillations offers an exciting opportunity, cortical gamma oscillations can be induced by multiple mechanisms (Tiesinga and Sejnowski, 2009; Buzsaki and Wang, 2012). While it is likely that different methods to induce cortical gamma oscillations (e.g., optogenetic activation or sensory stimulation) activate distinct neural ensembles, it is unclear whether other approaches to induce cortical gamma oscillations can also be beneficial. Addressing this issue is important to determine the underlying molecular and cellular mechanisms of how cortical gamma oscillations can modify AD pathology.

In the present study, we focus on the basal forebrain (BF) because of the following reasons: first, the BF is one of the most affected brain regions in AD (Whitehouse et al., 1982; Ballinger et al., 2016; Schmitz et al., 2016). Second, BF neurons provide cortex-wide projections (Zaborszky et al., 2012; Lin et al., 2015; Xu et al., 2015; Ballinger et al., 2016; Do et al., 2016; Gielow and Zaborszky, 2017). Third, the activation of BF neurons can modify cortical states (Kim et al., 2015; Sakata, 2016). Thus, the BF can be a unique target to modulate cortex-wide neural activity. While the BF consists of multiple nuclei and cell types (Zaborszky, 2002; Zaborszky et al., 2012; Xu et al., 2015; Ballinger et al., 2016), we are particularly interested in parvalbumin-positive (PV+) neurons (McKenna et al., 2013; Kim et al., 2015; Do et al., 2016).

BF PV+ neurons strongly innervate the pallidum, the striatum including the lateral septal complex, the anterior cingulate area, the infralimbic area, the retrosplenial area, the hippocampus, the thalamus, the lateral hypothalamus, the motor-related midbrain region and the behavioral state-related pontine region (Do et al., 2016). Gamma oscillations in the frontal cortex can be induced by optogenetic stimulation of BF PV+ neurons (Kim et al., 2015). To test the hypothesis that cortical gamma oscillations can modify AD pathology, we induced cortical gamma oscillations by optogenetically stimulating BF PV+ neurons in 5xFAD mice, the most aggressive AD mouse model (Oakley et al., 2006). Contrary to previous observations, we observed increased Aβ_1-42_ in the frontal cortical region, but not Aβ_1-40_. We also found that amyloid plaques increased in the medial prefrontal cortex and the septal nuclei. These results imply distinct molecular and cellular responses to different approaches to induce cortical gamma oscillations.

## Materials and Methods

### Animals

All animal experiments were performed in accordance with the United Kingdom Animals (Scientific Procedures) Act of 1986 Home Office regulations and approved by the Home Office (PPL70/8883). All transgenic lines were maintained on a C57BL/6 background. 5xFAD mice (JAX006554) (Oakley et al., 2006) were crossed with PV-IRES-Cre mice (JAX008069), followed by crossing with Ai32 mice (JAX012569). All genotyping was performed by Transnetyx using real-time PCR. Six 4-6 months old animals in both sexes were used in this study: three 5xFAD::PV-Cre::Ai32 mice (1 male, 2 females) and three 5xFAD::PV-Cre (2 males and 1 female). After surgery described below, they were housed individually in high-roofed cages with a 12 h:12 h light/dark cycle (light on hours: 7:00-19:00). They had ad libitum access to food and water. All experiments were performed during the light period.

### Surgery

Mice were anesthetized with isoflurane (5% for induction, 1-2% for maintenance) and placed in a stereotaxic apparatus (SR-5M-HT, Narishige). Body temperature was maintained at 37°C with a feedback temperature controller (50-7221-F, Harvard bioscience). Lidocaine (2%, 0.1 mL) was administered subcutaneously at the site of incision. Carprofen (Rimadyl, 5 mg/kg) was also administered subcutaneously at the back. After incision, the skull was exposed and cleaned. Four bone screws were implanted on the skull for monitoring cortical electroencephalograms (EEGs) (coordinate for frontal EEGs: AP, 1.5 mm; ML, ±1 mm) (coordinate for parietal EEGs: AP, −2 mm; ML, ±2 mm). An additional bone screw was implanted over the cerebellum as a ground/reference. These screws were connected to a two-by-three connector (SLD-112-T-12, Samtec). An optical fiber canula (CFM14L05, Thorlabs) was implanted to target the basal forebrain (AP, 0 mm; ML, 1.6 mm; DV, 5 mm from the cortical surface), including the substantia innominate, the magnocellular nucleus, and the diagonal band nucleus, and fixed with dental cement. After a recovery period (at least 5 days), mice were habituated to an open field (21.5 cm × 47 cm × 20 cm) by connecting a 16-channel amplifier board (RHD2132, Intan Technologies) and an interface cable as well as a patch cable (OPT/PC-FC-LCF-200/230-1.0L, Plexon).

### Electrophysiological recording and optogenetic stimulation

Electrophysiological signals were amplified relative to a cerebellar bone screw and were digitized at 1 kHz (RHD2132 and RHD2000, Intan Technologies). These signals were fed into a data acquisition device (NI-USB-6002, National Instruments), which was controlled by custom-made LabVIEW code running on a PC. The recording was begun by >10 min baseline recording (Pre), followed by 1-h optical stimulation (Stim). Pulses of blue light (470 nm, PlexBright, Plexon) were delivered 40 Hz for one hour through the patch cable attached to the implanted optical fiber. The light output at tip of the fiber was 114-178 mW/mm^2^. The recording was finished by another >10 min baseline recording (Post). The same procedure was repeated over three consecutive days.

### Tissue collection

After the final recording session, mice were deeply anesthetized with mixture of pentobarbital and lidocaine and perfused transcardially with saline. The brains were removed immediately and trimmed into two hemispheres. One of which was used for enzyme-linked immunosorbent assays (ELISA) whereas the remaining hemisphere was immersed overnight in 4% paraformaldehyde/0.1 M phosphate buffer, pH 7.4, at 4 °C for histological analysis. For ELISA, the prefrontal cortex and basal forebrain were isolated, immediately frozen and stored at −80°C until use.

### ELISA

Frozen tissue samples were homogenized in phosphate buffer saline (PBS) containing protease and phosphatase inhibitor cocktails (78440, ThermoFisher) and centrifuged at 2000 rpm for 5 min. The supernatants were then transferred to new tubes and centrifuged at 13,000 rpm for 15 min. The supernatants were collected and subjected to Aβ measurement with the use of mouse Aβ_1-40_ (KMB3481, ThermoFisher) or Aβ_1-40_ ELISA kit (KMB3441, ThermoFisher).

### Histology

The fixed brain tissue was immersed in a 30% sucrose in PBS at 4 °C. The brains were frozen and were cut into coronal sections with a sliding microtome (SM2010R, Leica) with a thickness of 50 μm. Sections were stained for either PV or plaques. For PV staining, sections were washed with PBS for 5 min, three times at room temperature (RT) and then incubated in a blocking solution (10% normal goat serum, NGS, in 0.3% Triton-X in PBS, PBST) for 1 h at RT followed by incubating with primary antibodies (anti-PV, 1:1000-2000, P3088, Sigma-Aldrich) in 3% NGS in PBST at 4 °C overnight. After washing, sections were incubated with secondary antibodies (goat anti-mouse Alexa Fluor 594, 1:1000, A-11005, ThermoFisher) for 2 h at RT. For plaques staining, sections were washed with PBS for 5 min, three times at RT and then incubated with Thiazine-Red (0.05%, cat#27419.123, VWR) for 20 min. After washing, sections were counter-stained with DAPI (1:1000), mounted on gelatin-coated slides and cover-slipped. Epifluorescence images were captured at 4x magnification with a digital CMOS camera (C11440-36U, Hamamatsu) using freely available software (WinFluor). Captured brain regions were the anterior cingulate area (ACA), the medial prefrontal cortex (mPFC) including the prelimbic and infralimbic areas, the primary somatosensory cortex (S1), the entorhinal cortex (ENT), the dentate gyrus (DG), the subiculum (SUB), the septal nuclei (SEP) including both the lateral and medial septum, and the ventral posteromedial nucleus of the thalamus (TH).

### Data analysis

For spectral analysis of EEGs, MATLAB (version 2018b, Mathworks) was used with Signal Processing Toolbox and Chronux Toolbox (http://chronux.org/). To assess the changes in spectral power at a certain frequency band, power spectral density (PSD) for each period (Pre, Stim, or Post period) was estimated with Welch’s method and the sum of PSD at the frequency band was computed. The summed PSD was further normalized by the summed PSD in Pre period for comparisons.

For image analysis, a region of interest for each brain region (see above) was manually determined with Fiji and processed with a custom-written MATLAB code. The number of plaques was automatically quantified as follows: first, each image was binarized by computing a threshold, which was defined as *M* + 2.8 × *D*, where *M* is the median of pixel intensities and *D* is the average absolute deviation of pixel intensities. After applying a median filter, detected particles with a pre-determined range of sizes were recognized as plaques. Finally, the density of plaques was computed.

### Statistics

Data was presented as mean ± SEM. Statistical analyses were performed with MATLAB. Student’s *t*-test was performed in **Figure 2A**. One-way ANOVA and two-way ANOVA with post-hoc Tukey’s Honest Significant Difference (HSD) test were performed in **Figures 1E and 2D**, respectively.

**Figure 1.**
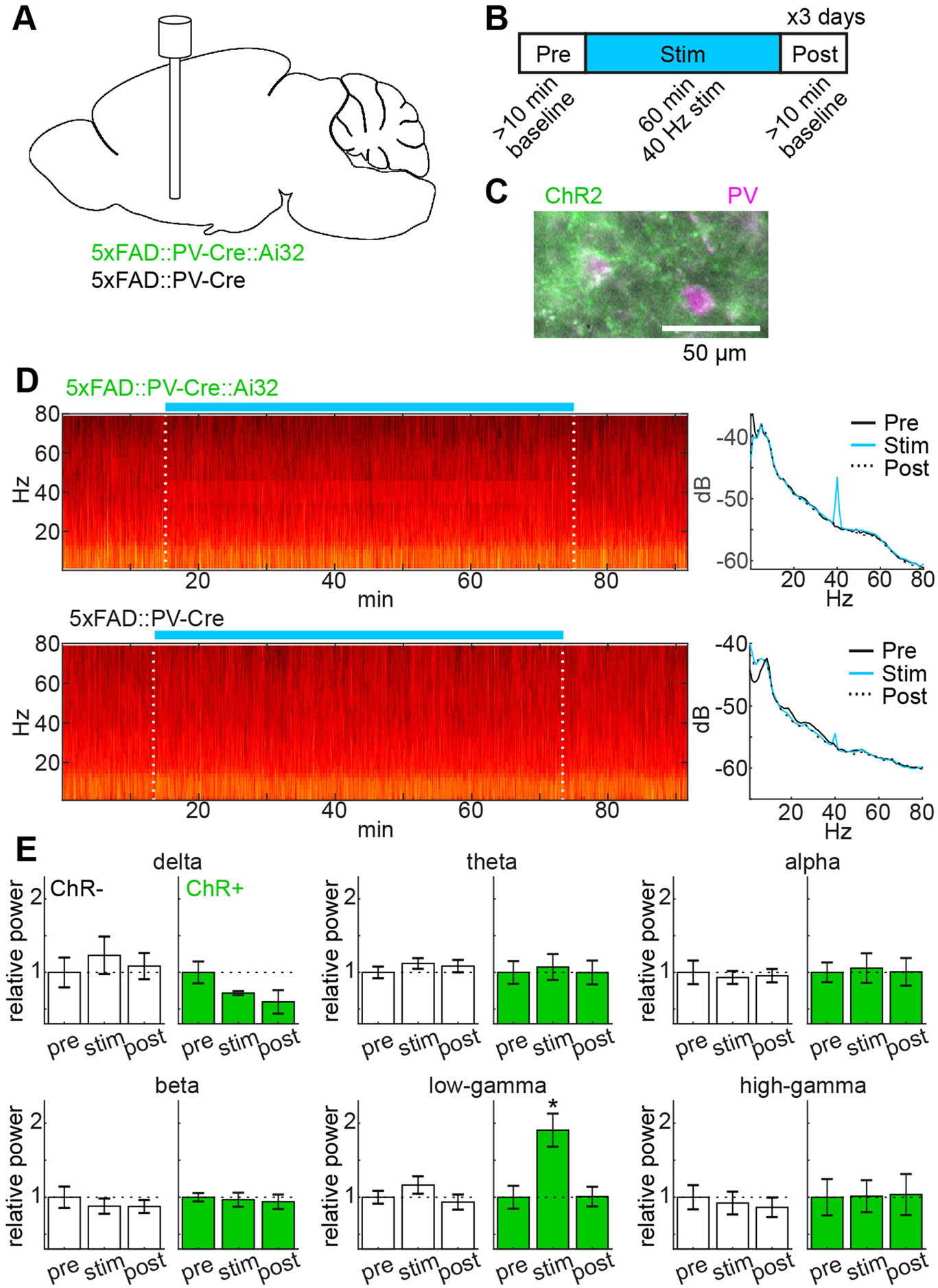
The induction of cortical gamma oscillations by optogenetic stimulation of BF PV+ neurons in 5xFAD mice. (**A**) A diagram of the experimental approach, showing the optical fiber implant in the BF and genotypes used in this study. Ai32 is a Cre-dependent ChR2 mouse. (**B**) A timeframe of the optogenetic experiment. (**C**) A photograph, showing co-expression of ChR2-EYFP and PV in the BF. (**D**) Examples of spectrogram (*left*) and power spectral density (*right*) in ChR2+ (*top*) and ChR2− (*bottom*) animals. (**E**) Relative changes in spectral power across frequency bands in ChR2+ (*green*) and ChR2− groups (*white*). delta, 0.5-4 Hz; theta, 5-8 Hz; alpha, 8-12 Hz; beta,15-30 Hz; low gamma, 38-43 Hz; and high gamma, 50-80 Hz. *, *p* < 0.05 (one-way ANOVA with post-hoc HSD test).

**Figure 2.**
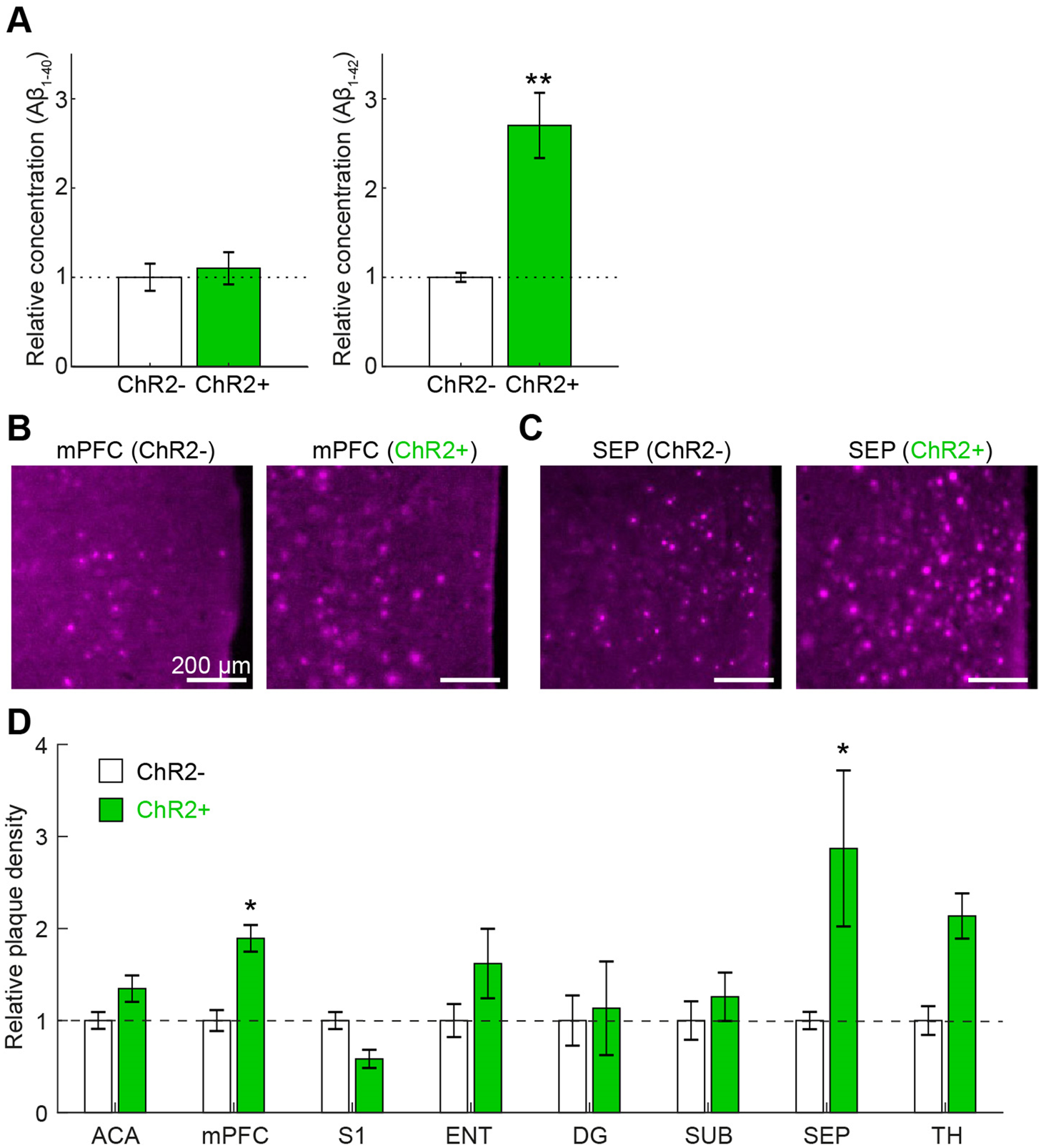
Biochemical and histological analysis of changes in amyloid load following optogenetic stimulation. (**A**) Relative concentration of Aβ_1-40_ (*left*) and Aβ_1-42_ (*right*) in the prefrontal cortical area, measured by ELISA. **, *p* < 0.01 (*t*-test). (**B and C**) Photographs of thiazine-red-stained sections in the mPFC (**B**) and the SEP (**C**). (**D**) Plaque density across brain regions. The values in each brain region were normalized by the average plaque density in ChR2− group. *p* < 0.05, two-way ANOVA with post-hoc HSD test. ACA, anterior cingulate area; mPFC, medial prefrontal cortex; S1, primary somatosensory cortex; ENT, entorhinal cortex; DG, dentate gyrus; SUB, subiculum; SEP, septum; TH, ventral posteromedial nucleus of the thalamus.

## Results

### The induction of cortical gamma oscillations by optogenetic stimulation of basal forebrain PV+ neurons

To induce cortical gamma oscillations, we expressed ChR2 in PV+ neurons of 5xFAD mice (**Figs. 1A and C**) and applied 40 Hz optical stimulation for 1 hour (**Fig. 1B**) in the basal forebrain (BF), including the substantia innominate, the magnocellular nucleus, and the diagonal band nucleus. As previously reported (Kim et al., 2015), 40 Hz optical stimulation elicited the strong modulation in cortical EEGs at 40 Hz (**Fig. 1D**). To quantify this tendency, we computed spectral power in each period (Pre, Stim, and Post periods) across the following frequency bands: delta (0.5-4 Hz), theta (5-8 Hz), alpha (8-12 Hz), beta (15-30 Hz), low gamma (38-43 Hz), and high gamma (50-80 Hz) (**Fig. 1E**). Although we observed a tendency of reduction in the delta power, the effect was not significant (*F*_2,6_ = 2.63, *p* = 0.15, one-way ANOVA, *n* = 3 animals in each condition). This tendency may be explained by the fact that activation of BF PV+ neurons leads to arousal (Kim et al., 2015; Xu et al., 2015). We confirmed that the low gamma power increased only during optical stimulation period in ChR2 expressing animals (*F*_2,8_ = 8.91, *p* = 0.043, one-way ANOVA with post-hoc HSD test, *n* = 3 animals in each condition). Thus, optogenetic stimulation of BF PV+ neurons can induce cortical gamma oscillations in 5xFAD mice.

### Increased amyloid burden following optogenetic stimulation

To examine whether optogenetic stimulation affects AD pathology, we collected brain samples after three-day optogenetic stimulations and measured the concentration of Aβ_1-40_ and Aβ_1-42_ by performing ELISA (**Fig. 2A**). Because BF PV+ neurons project primarily to the prefrontal cortical regions (Do et al., 2016), we analyzed amyloid load in the prefrontal cortical area and the BF separately. Compared to ChR2-group, we found significant increase in the concentration of Aβ_1-42_ in the prefrontal cortical area (*p* = 0.0099, *t*-test, *n* = 3 animals in each condition) and the BF (1.00 ± 0.12 in ChR2-, 2.4 ± 0.48 in ChR2+, *p* = 0.047, *t*-test, *n* = 3 animals in each condition) whereas we did not find any significant changes in prefrontal Aβ_1-40_ (*p* = 0.69, *t*-test, *n* = 3 animals in each condition) and BF Aβ_1-40_ (1.00 ± 0.02 in ChR2-, 1.02. ± 0.06 in ChR2+, *p* = 0.74, *t*-test, *n* = 3 animals in each condition).

To verify these observations, we also performed histological analysis across multiple brain regions (**Figs. 2B-D**). We analyzed the anterior cingulate area (ACA) (*n* = 16 sections of ChR2-from 3 animals, *n* = 18 sections of ChR2+ from 3 animals), the medial prefrontal cortex (mPFC) (*n* = 16 vs *n* = 11), the primary somatosensory cortex (S1) (*n* = 8 vs *n* = 5), the entorhinal cortex (ENT) (*n* = 7 vs *n* = 4), the dentate gyrus (DG) (*n* = 6 vs *n* = 7), the subiculum (SUB) (*n* = 6 vs *n* = 7), the septum (SEP) (*n* = 8 vs *n* = 7), and the ventral posteromedial nucleus of the thalamus (TH) (*n* = 7 vs *n* = 6). We found that the mPFC (**Fig. 2B**) and the SEP (**Fig. 2C**) exhibit significant increase in the number of amyloid plaques in ChR2+ groups (*F*_7,123_ = 2.60, *p* < 0.05, two-way ANOVA with post-hoc HSD test) (**Fig. 2D**). These results indicate that the induction of cortical gamma oscillations by optogenetically stimulating BF PV+ neurons increases amyloid load in several brain regions.

## Discussion

The induction of cortical gamma oscillations can modify AD pathology. Here we found that optogenetic activation of BF PV+ neurons in 5xFAD mice increases amyloid load in several brain regions, including the mPFC and the septal nuclei. Our results suggest that effects of cortical gamma oscillations on AD pathology can depend on their induction mechanisms.

Previous studies showed that optogenetic activation of BF PV+ neurons can induce cortical gamma oscillations and arousal (Kim et al., 2015; Xu et al., 2015). We confirmed the optogenetic induction of gamma oscillations in 5xFAD mice (**Fig. 1**). Although we initially concerned that the sharp increase in low gamma power might have been due to an optical artifact, we did not observe a similar modulation in the control group, indicating that the increase in low gamma power can be explained by the expression of ChR2. We also observed a tendency of the reduction in delta power. This observation is also consistent with the notion that the activation of BF PV+ neurons induce arousal (Xu et al., 2015).

While BF PV+ neurons project to a wide range of brain regions, the mPFC and the septum are also target regions (Do et al., 2016). Our results are consistent with this anatomical observation in sense that optogenetic activation of BF PV+ neurons can modify their downstream neurons. However, we did not directly monitor neural activity of BF PV+ neurons and the spatial extent of optical stimulation effects. This issue remains to be addressed. With respect to the spatial extent of optical stimulation effects, because we took a bi-genetic approach where a Cre-dependent optogenetic mouse line (i.e., Ai32) was crossed with PV-IRES-Cre mice, most of PV+ neurons across the brain are supposed to express ChR2. In future, it would be worth confirming our results by adopting a viral approach to express opsins only in the BF.

The increase in amyloid load (**Fig. 2**) was totally unexpected because previous studies show that the induction of cortical gamma oscillations can reduce amyloid burden and improve cognitive functions (Iaccarino et al., 2016; Adaikkan et al., 2019; Martorell et al., 2019). What can explain this discrepancy? The method to induce cortical gamma oscillations is different at least. Whereas previous studies induced cortical gamma oscillations by either optogenetic stimulation of cortical PV+ neurons or 40 Hz sensory stimulus, we induced cortical gamma oscillations by activating BF PV+ neurons. Because BF GABAergic neurons preferentially target cortical interneurons (Freund and Meskenaite, 1992), our approach could suppress cortical PV+ neurons, rather than activating them. Because cortical gamma oscillations can be induced by multiple mechanisms (Tiesinga and Sejnowski, 2009; Buzsaki and Wang, 2012), it is possible that these methodological differences may induce distinct molecular and cellular responses, resulting in opposing effects on AD pathology. In addition, because microglia plays a key role in AD pathogenesis (Hickman et al., 2018; Henstridge et al., 2019; Long and Holtzman, 2019) and microglia exhibits highly dynamic responses depending on global brain states (Liu et al., 2019; Stowell et al., 2019), it is also conceivable that the optogenetic activation of BF PV+ neurons induces not just cortical gamma oscillations, but also changes in neuromodulatory tones, leading to microglial responses in the cortex different from other approaches.

Although our finding regarding the modification of Aβ_1-42_ (**Fig. 2A**) is consistent with another optogenetic study (Yamamoto et al., 2015), it remains to be determined how Aβ_1-42_ was selectively increased. While both forms of Aβs differ in their metabolism and aggregation mechanisms (Qiu et al., 2015) and the cerebrospinal fluid Aβ_1-42_/Aβ_1-40_ ratio has been considered as a diagnostic marker of AD (Janelidze et al., 2016), further investigation on activity-dependent regulation of Aβs will provide insight into the pathogenesis of AD.

In summary, optogenetic activation of BF PV+ neurons in 5xFAD mice increases amyloid load. Although the induction of cortical gamma oscillations is still a promising approach to modify AD pathology, the outcomes can vary depending on how to induce gamma oscillations. A better understanding of underlying circuit mechanisms of gamma oscillations as well as molecular and cellular responses to distinct gamma oscillations is crucial to fully implement this novel approach for treatment and prevention of AD and other neurodegenerative diseases.

## Acknowledgements

We thank Mark Barbour for helping ELISA. This work was supported by BBSRC (BB/M00905X/1) and Alzheimer’s Research UK (ARUK-PPG2017B-005) to SS.

## Authors contribution

CAW and SS designed and conceived the project. CAW and SS performed experiments. SF and SS analyzed data. SS wrote the manuscript with inputs from CAW and SF.

